# miR-223 Plays A Critical Role in Obesogen-Enhanced Adipogenesis in Mesenchymal Stem Cells and in Transgenerational Obesity

**DOI:** 10.1101/2022.10.27.514142

**Authors:** Richard C. Chang, Erika M. Joloya, Zhuorui Li, Bassem M. Shoucri, Toshi Shioda, Bruce Blumberg

## Abstract

Exposure of pregnant F0 mouse dams to the obesogen tributyltin (TBT) predisposes unexposed male descendants to obesity and diverts mesenchymal stem cells (MSCs) toward the adipocytic lineage. TBT also promotes adipogenic commitment and differentiation of MSCs, in vitro. We sought to identify TBT-induced factors predisposing MSCs toward the adipocytic fate. We exposed mouse MSCs to TBT, the PPARγ-selective agonist rosiglitazone or the RXR-selective agonist LG-100268 and determined their transcriptomal profiles to determine candidate microRNAs (miR) regulating adipogenic commitment and differentiation. Of the top 10 candidate microRNAs predicted by Ingenuity Pathway Analysis, miR-21, miR-33 and miR-223 were expressed in a manner consistent with an ability to differentially regulate target genes during adipogenesis. After 24 hours exposure to 50 nM TBT, miR-223 levels in MSCs were increased and expression of its target genes ZEB1, NFIB, and FOXP1 was decreased. Both ROSI and TBT increased miR-223 levels, and this induction was inhibited by the PPARγ antagonist T0070907 but not by the RXR antagonists HX531 or UVI3003, placing miR-223 downstream of PPARγ. Chromatin immunoprecipitation confirmed TBT-induced binding of PPARγ to regulatory elements in the miR-223 promoter. miR-223 levels were elevated in white adipose tissue of F2 and F3 male descendants of pregnant F0 mouse dams exposed to 50 nM TBT throughout gestation. miR-223 levels were further induced in males fed with an increased fat diet. We infer that TBT induced miR-223 expression and increased adipogenesis in MSCs through the PPARγ pathway and that transgenerationally increased expression of miR-223 plays an important role in the development of obesity caused by TBT exposure.

## Introduction

Obesity is a worldwide public health issue that, although multifactorial in nature, is generally considered to be the result of energy imbalance [1]. Compelling evidence suggests that obesity is a critical contributor to the development of adipose tissue inflammation and insulin resistance [2, 3], two major causal factors in the pathogenesis of obesity-associated type 2 diabetes [4]. Therefore, adipose tissue itself is believed to play an important role in regulating obesity-associated health issues.

A growing number of studies have linked exposure to endocrine-disrupting chemicals (EDCs) and the obesity pandemic [5–8]. One subset of EDCs, obesogens, has been found to increase white adipocyte number and/or size and consequently promote adiposity [9]. The obesogens tributyltin (TBT) and triphenyltin (TPT) activated the peroxisome proliferator activated receptor gamma (PPARγ), the master regulator of adipogenesis, and its heterodimeric partner, the retinoid X receptor (RXR) [10–13]. In vitro studies showed that TBT promoted adipogenesis at low nanomolar levels in both human and mouse multipotent mesenchymal stromal stem cells (a.k.a., mesenchymal stem cells, MSCs) [14, 15]. Using a mouse model, we showed that exposure of pregnant F0 mouse dams to TBT predisposed MSCs towards the adipose lineage in F1 offspring [15] and increased lipid accumulation in liver and white adipose depots in F1, F2, and F3 generations [9]. Interestingly, male F4 descendants of F0 dams treated with TBT throughout pregnancy [16] or pregnancy and lactation [17] exhibited a large increase in fat storage compared with controls when fed with a diet containing modestly increased fat content (from 13.2% to 21.6% Kcal from fat).

MicroRNAs (miRs) are a family of small non-coding RNAs of 21-23 nucleotides in length. miRs typically regulate protein synthesis at the post-transcriptional level by targeting specific mRNAs through perfect or imperfect base-pairing and triggering mRNA degradation or translational repression, respectively [18]. Several studies suggested possible roles for miRs in adipogenesis. By targeting adipogenic transcriptional factors, such as PPARγ and CCAAT-enhancer-binding proteins (C/EBPs), miR-155 and miR130 decreased white adipocyte formation [19, 20]. On the other hand, miR-210 targeted TCF7l2 in the Wnt/β-catenin signaling pathway and promoted adipogenesis [21].

miR-223 is a highly conserved microRNA that was first reported to play a role in myeloid differentiation and inflammation [22, 23]. miR223 levels were reduced in hepatocellular carcinomas [24] and leukemias [25, 26] while its overexpression reduced viability of cancer cells [24]. miR-223 regulated polarization of adipose-resident macrophages by targeting PBX/Knotted 1 Homeobox 1 (Pknox1). This resulted in suppression of nuclear factor kappa B (NF-κB) and c-Jun N-terminal kinase (JNK) pathways and enhanced adipocyte response to insulin stimulation [27, 28]. Increased miR-223 expression was also reported during adipogenesis in mouse bone marrow stromal cells [29]. Here, we show that the obesogen TBT induces miR-223 expression through the PPARγ pathway in mouse MSCs, leading to, and enhancing adipogenesis. We also provide evidence that treatment of pregnant F0 mouse dams with TBT led to increased miR-223 expression in the white adipose tissue (WAT) of male descendants through the F3 generation and that miR-223 expression is further elevated when dietary fat is increased. These observations support an important role for miR-223 downstream of PPARγ in the transgenerationally inherited predisposition to obesity.

## Materials and Methods

### Chemicals and Reagents

TBT, LG100268, HX531, UVI3003, dexamethasone, isobutylmethylxanthine, insulin, Nile Red, and Hoechst 33342 were purchased from Sigma-Aldrich (St. Louis, MO). T0070907 was purchased from Enzo Life Sciences (Farmingdale, NY). Rosiglitazone (ROSI) was purchased from Cayman Chemical (Ann Arbor, MI).

### Cell Culture

Bone marrow derived multipotent MSCs from the long bones of C57BL/6J mice were purchased at passage 6 (OriCell; Cyagen Biosciences, CA) and stored at passage 8 or 9 in liquid N2. Cells were maintained, as previously described [9], in Dulbecco’s modified Eagle medium (DMEM) containing 10% calf bovine serum, 10 mM HEPES, 1 mM sodium pyruvate, 100 IU/mL penicillin, and 100 μg/mL streptomycin [13]. MSCs were plated at 60,000 cells/cm^2^ in 12-well cell culture plates for adipogenesis assays. Cells were allowed to attach and acclimate for 24 hours prior to 48 hours of chemical treatment or adipogenesis. Specific ligands, 100 nM ROSI, 50 nM TBT, and 200 nM LG268 were dissolved in DMSO and administered every 3 days throughout the duration of the adipogenesis assay. Antagonist assays were performed in parallel under the same conditions. The antagonist T0070907 (50nM), HX531 (100nM), UVI3003 (500nM) or DMSO vehicle control was added every 12 hours. The amount of dimethyl sulfoxide (DMSO) vehicle was kept at < 0.1% in all assays. miRCURY ™LNA microRNA inhibitor (anti-miR-223) (Qiagen, Hilden, Germany) is the sequence-specific and chemically modified oligonucleotide that specifically target and knockdown miR-223 miRNA molecules.

### Adipogenesis assay

Once cells reached 100% confluency in culture plates, cells were induced to differentiate with an adipose induction cocktail (500 μM isobutylmethylxanthine, 1 μM dexamethasone, and 5 μg/mL insulin) in minimal essential medium α (αMEM) containing 15% fetal bovine serum, 10 mM HEPES, 2 mM l-glutamine, 100 IU/mL penicillin, and 100 μg/mL streptomycin as we described [17]. Cells were replenished with fresh media, differentiation factors, and chemical ligands every 3 days. Cells were differentiated over the course of 14 days, fixed in buffered 3.7% formaldehyde, followed by one wash of PBS for 1 minute, and then maintained at 4°C in PBS overnight to remove residual phenol red. To quantify lipid accumulation, Nile Red (1 μg/mL) was used to stain neutral lipids and Hoechst 33342 (1 μg/mL) to stain DNA. For each biological replicate, Nile Red fluorescence units (excitation 485 nm, emission 590 nm) (RFU) were measured relative to Hoechst RFU (excitation 355 nm, emission 460 nm) using a SpectraMax Gemini XS spectrofluorometer (Molecular Devices, Sunnyvale, CA) using SoftMax Pro (Molecular Devices) [13].

### qPCR

Differentiated adipocytes were lysed with Trizol following the manufacturer’s recommended protocol (Thermo Fisher Scientific, MA) and total RNA recovered after isopropanol precipitation (Fisher Chemical, PA). Complementary DNA of microRNAs was synthesized from 5 μg total RNA using microRNA-specific primers (Qiagen, Hilden, Germany) according to the manufacturer’s instructions. Gene expression was assessed with real-time quantitative polymerase chain reaction (qPCR) using SYBR_™_ Green PCR Master Mix (Thermo Fisher Scientific, MA) on a Roche LightCycler 480 II (Roche, Switzerland). Cycle threshold values were quantified as the second derivative maximum using LightCycler software (Roche, Switzerland). Noncoding small RNA control sno202 served as an endogenous reference gene [30]. The 2^-ΔΔCt^ method [31] was used to analyze RT-qPCR data and determine relative quantification. Standard propagation of error was used throughout for each treatment group [32, 33]. Error bars represent the SEM from three to four biological replicates, calculated using standard propagation of error.

### Chromatin immunoprecipitation

Chromatin immunoprecipitation (ChIP) was performed using an established method as previously described [13]. Briefly, MSCs were plated and treated in 10-cm dishes (Thermo-Nuncalon) to obtain adequate material. At the end of chemical treatment (48 hours), cells were fixed at room temperature for 10 minutes with 1% paraformaldehyde (Fisher Chemical, PA) in DMEM, followed by an ice-cold phosphate-buffered saline wash, and then quenched for 5 minutes with 125 mM glycine at room temperature. Fixed cells were washed and then collected by scraping the plates, centrifuging, and then resuspending in phosphate-buffered saline at 10^7^ cells/mL. Equal numbers of cells were snap-frozen in liquid N2 and stored at −80°C. To isolate nuclei, cell pellets were lysed at 4°C for 10 minutes with a gentle detergent recipe consisting of 50 mM HEPES-KOH, pH 7.5, 140 mM NaCl, 1 mM EDTA, 10% glycerol, 0.5% Nonidet P-40, 0.25% Triton X-100, Halt™ Protease Inhibitor Cocktail (Thermo Fisher Scientific, MA). Next, nuclei were recovered by centrifugation at 8000xg for 15 minutes, washed for 10 minutes at room temperature (10 mM Tris-HCl, pH 8.0, 200 mM NaCl, 1 mM EDTA, 0.5 mM EGTA, protease inhibitors (Thermo Fisher Scientific, MA)), and lysed in 300μL nuclear lysis buffer (10 mM Tris-HCl, pH 8.0, 200 mM NaCl, 1 mM EDTA, 0.5 mM EGTA, 0.1% Na-deoxycholate, 0.5% N-lauroylsarcosine, protease inhibitors (Thermo Fisher Scientific, MA)). Chromatin samples were prepared by sonicating in 0.5 mL thin-walled polymerase chain reaction tubes (BrandTech, CT) using a QSonica Q800R2 (QSonica, CT) with the following settings: 30 seconds on/30 seconds off, amplitude 40% repeated for 30 minutes. Triton X-100 (1%) was added to sonicated lysates prior to high-speed, cold centrifugation to remove debris. A total of 5μg DNA was immunoprecipitated with preblocked protein A/G Dynabeads (Thermo Fisher Scientific, MA) complexed to 2.5 μg antibody (anti-PPARγ, ab233218, Abcam, Cambridge, UK;). Beads were washed three times with LiCl buffer (50 mM HEPES-KOH, pH 7.5, 500 mM LiCl, 1 mM EDTA, 1% Nonidet P-40, 0.7% Na-deoxycholate) and once with Tris-EDTA buffer plus 50mM NaCl. To release chromatin from beads, pelleted beads were resuspended in elution buffer (50mM Tris-HCl, pH 8.0, 10 mM EDTA, 1% sodium dodecyl sulfate) and incubated at 65°C for 30 minutes. Cross-link reversal was performed overnight at 65°C. DNA samples were purified using the ChIP DNA Clean & Concentrator kit (Zymo Research, CA) following RNase A (0.2 mg/mL, 2 hours, 37°C) and proteinase K (0.2 mg/mL, 2 hours, 55°C) treatment. Input DNA content was determined by spectrophotometry (Nanodrop, Thermo Fisher Scientific, MA).

### RNA-sequencing and Bioinformatics

Transcriptomal profiles were generated and published previously [13]. In brief, total RNAs integrity was verified using Tapestation high-sensitivity RNA screen tapes (Agilent Technologies, Santa Clara, CA), RNA integrity numbers ranged from 8.0 to 8.6. Strand-specific, barcode-indexed RNA sequencing (RNA-seq) deep-sequencing libraries were synthesized from total RNA with ERCC spike-in controls (Thermo Fisher Scientific, MA) using Ovation RNA-Seq Systems 1-16 for Mouse (NuGen Technologies, CA). Tapestation analysis determined the size distribution of the libraries to be 200 to 800 bp, peaking at 300 bp. KAPA Illumina library quantification kit (KAPA Biosystems, MA) was utilized to quantify RNA-seq libraries, and up to 12 libraries were multiplexed in each run of the Illumina NextSeq500 deep sequencer (75 nt + 75 nt, paired-end) to generate fastq raw read sequence files which were aligned to the mouse genome reference sequence GRCm38/mm10 using the STAR aligner [34], and the resulting bam format aligned reads were subjected to QC analysis using fastQC (Babraham Institute, Cambridge, United Kingdom), followed by extraction of uniquely mapped reads using SAMtools [35]. Differential expression was assessed in DESeq2 [36] using the DESeq function with α = 0.01. Differentially expressed genes (DEGs) were defined by Benjamini–Hochberg corrected *p* values (p-adj < 0.01). Sequencing files have been uploaded and are available on Gene Expression Omnibus (GSE216429). Candidate microRNA enrichments were computed by the microRNA-mRNA correlation feature supported by Ingenuity Pathway Analysis (IPA) [37]. P values were corrected for multiple testing using the Benjamini–Hochberg method.

### Mouse treatment

We undertook a new transgenerational experiment based on our previous transgenerational model[17], denoted as T4. Briefly, 148 female and 50 male C57BL/6J mice (5 weeks of age) were purchased from The Jackson Laboratory and randomly assigned to treatment groups and exposed via drinking water to 50 nM TBT or 0.1% DMSO vehicle (all diluted in 0.5% carboxymethyl cellulose in water to maximize solubility) for 7 days prior to mating as we have described [16]. Sires were never exposed to the treatment. Treatment was removed during mating, then resumed after males were removed until F1 (1^st^ generation) offspring were born. F2 (2^nd^ generation) offspring were exposed to TBT as germ cells in the exposed F1 embryos. F3 (3^rd^ generation) offspring were not exposed to TBT. One male and one female were randomly selected per litter for analysis. In diet challenge experiment, designated F2 (14 males and 14 females for each group) and F3 (15 males and 15 females for each group) were switched to a higher fat diet (PicoLab Rodent Chow, 5058, 21.4% Kcal from fat) whereas control groups (F2: 15 males and 15 females; F3: 12 males and 12 females) were maintained on a standard chow diet (PicoLab Rodent Chow, 5053, 13.2% Kcal from fat). F2 offspring started diet challenge at 4 weeks age for 8 weeks when a significant fat content increase was confirmed and persisted. F3 offspring started diet challenge at week 17 for 5 weeks when a significant fat content increase was confirmed and persisted. Mice were fasted for 12 hours prior to euthanasia and tissue collection.

### Statistical Analysis

GraphPad Prism 7.0 (GraphPad Software, Inc.) was used to perform statistical analysis for all datasets. A one-way analysis of variance (ANOVA) followed by Dunnett’s post-hoc test was performed to compare the treatment group, ROSI, TBT, or LG268, to DMSO control in adipogenesis assays. In antagonist assays, treatment groups without antagonists were compared to a corresponding group co-treated with T0070907, HX531, or UVI3003 using a student’s t-test. *p* ≤ 0.05 was considered statistically significant.

## Results

### Expression of miR-21, miR-33, and miR-223 was significantly changed during adipogenesis of MSCs

We previously analyzed the transcriptomal profiles of MSCs during adipogenesis [13]. To predict candidate microRNAs involved in the 14-day process of adipose lineage commitment and differentiation in MSCs, we first utilized Ingenuity Pathway Analysis of differentially expressed genes defined by Benjamini-Hochberg corrected P values (p-adj < 0.01). The top ten predicted microRNAs were miR-124, miR-30, miR-29, miR-33, miR-8, miR-21, miR-515, miR-223, miR-16, and miR-221 (Figure S1). We next validated expression of these 10 microRNAs using real-time qPCR and found that 7 of them did not exhibit statistically significant changes in expression during adipogenesis (Figure S2) and were not considered further. miR-21 levels remained unchanged during adipogenesis in vehicle-treated MSCs whereas levels showed a decreasing trend that never reached statistical significance starting at adipogenesis day 4 in MSCs treated with 50 nM TBT (Figure 1A). miR-33 levels decreased during the 14-day adipogenesis in vehicle-treated MSCs, and TBT treatment partly relieved this suppression (Figure 1B). miR-223 levels increased by adipogenesis day 7 (2.62 fold compared to day 0) the continued to increase at day 10 (3.5 fold) and day 14 (5.35 fold) compared to day 0) (Figure 1C). This echoed previous studies of miR-223 expression in obese mice or humans [29, 38]. TBT treatment enhanced adipogenesis-induced miR-223 expression from 5.35 fold (DMSO) to 15.84 fold (TBT) by day 14 (Figure 1C). We next examined the transcriptomal profiles which were used to predict candidate microRNAs and confirmed that mRNA levels of three known miR-223 targets – namely, Foxp1, Zeb1, and Nfib – were significantly suppressed by both 50 nM and 100 nM TBT at days 4, 7, and 14 of adipogenesis (Figure 1D-F). We infer that TBT treatment accelerated adipogenesis-induced expression of miR-223, which inhibited expression of these target mRNAs.

**Figure 1.**
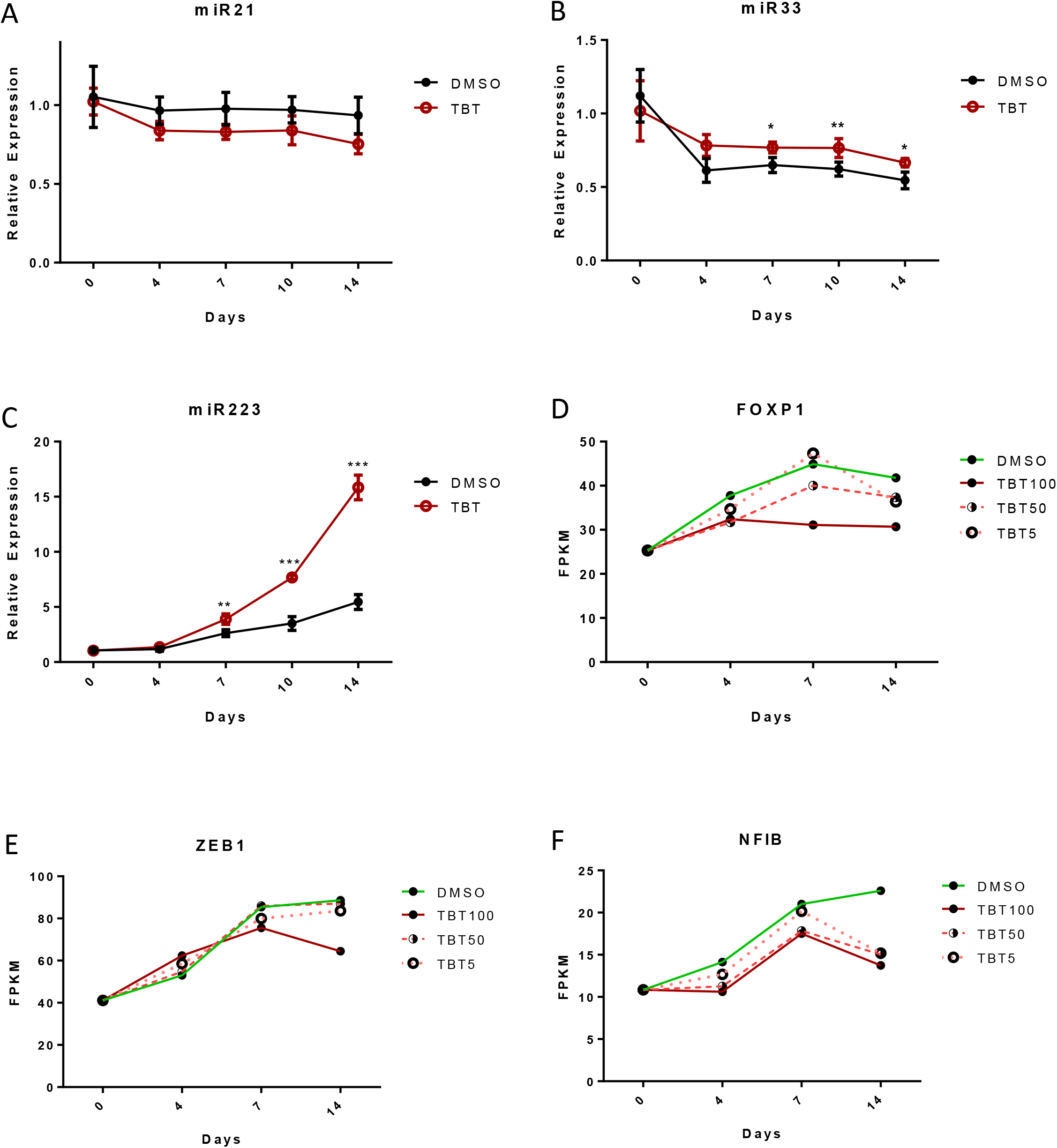
miR-223 expression is induced whereas mi-R223 target genes are suppressed during TBT-enhanced adipogenesis in MSCs. (A-C) qRT-PCR quantitation of three microRNAs, miR-21 (A), miR33 (B), and miR223 (C) in MSCs at adipogenesis day 0 −14. Expression was normalized to sno202 and presented as relative expression to vehicle control at day 0 (mean ± SEM, *P < 0.05, **P < 0.01 and ***P < 0.001). (D-F) RNA-seq determination of mRNA transcripts of miR-223 target genes FOXP1 (D), ZEB1 (E), and NFIB (F) in mMSCs exposed to 0-100 nM TBT at adipogenesis day 0-14. RNA-seq data were described in our previous study [52].

### TBT-induced miR223 expression is critical for TBT-enhanced adipogenesis

TBT acts through the nuclear receptor PPARγ and its heterodimeric partner RXR to promote adipogenesis in vitro and in vivo [9, 11, 13]. To elucidate the role of TBT-induced miR-223 in adipogenesis, we determined effects of 50 nM TBT on adipogenesis of MSCs as described [13]. The PPARγ-selective activator ROSI (100 nM) and the selective RXR activator LG268 (100 nM) were used as positive controls for PPARγ or RXR activation, respectively. microRNA suppression assays employed miR-223 antisense; a random antisense microRNA served as negative control. Transfection with the miR-223 antisense RNA significantly attenuated ROSI-induced and TBT-induced lipid accumulation after 14-days of adipogenesis (Figure 2A). Expression of Zeb1, Nfib, and Foxp1 was decreased by ROSI and TBT after 14-days of adipogenesis while co-treatment with miR-223 antisense reversed these effects (Figure 2B-D). We infer that TBT enhances miR-223 expression in MSCs to support in vitro adipogenesis and reduce steady-state levels of three target mRNA transcripts of miR-223. These data also support a role for PPARγ in regulation of miR-223 expression.

**Figure 2.**
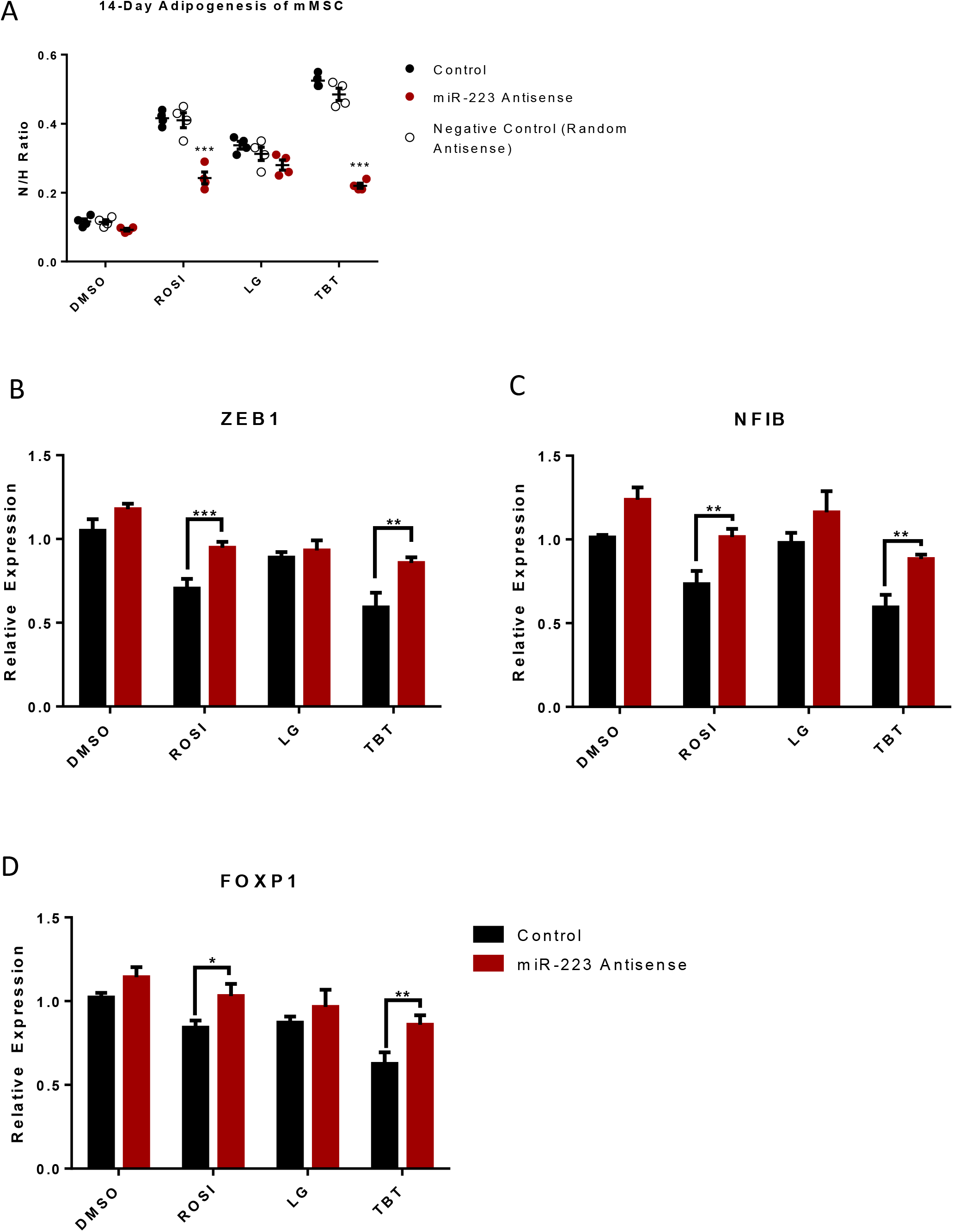
TBT-induced miR-223 expression contributes to TBT-enhanced adipogenesis in MSCs. (A) Adipogenesis assay was performed using mMSCs in the presence of adipogenesis cocktail (MDI). Cells were treated with a 100 nM ROSI, 50 nM TBT, or 100nM LG268 in presence of miR-223 antisense or random antisense as a negative control. All treatment groups were statistically compared to the control (black bars, 0.1% DMSO during 14 day adipogenesis). Lipid accumulation is shown as the ratio between fluorescence units (RFU) of Nile Red (N) and Hoechst (H), which were used to quantify lipid content and the number of cells per well, respectively. Each bar represents average of 4 replicates ± SEM. (B-D) qRT-PCR quantitation of mRNA transcripts of miR-223 target genes ZEB1 (B), NFIB (C), and FOXP1 (D) in mMSC-derived differentiated adipocytes at day 14. Expression was normalized to housekeeping gene, Ribosomal Protein Lateral Stalk Subunit P0 (RPLP0) and presented as relative expression to the group of DMSO exposure with negative control, the random antisense oligonucleotides (mean ± SEM, *P < 0.05, **P < 0.01 and ***P < 0.001).

### TBTinduced miR223 expression through PPARγ

To assess whether TBT enhanced miR-223 expression via RXR or PPARγ activation, we first asked if TBT alone induces miR-223 expression. We treated naïve MSCs with 50 nM TBT, 100 nM ROSI, or 100 nM LG268 for 24 or 48 hours (Figure 3). Expression of miR-21 and miR-33 expression was not changed by TBT (Fig. S3). Expression of miR-223 was increased in 24 hours by TBT (3.25 fold), ROSI (3.8 fold) and LG268 (2 fold) (Fig. 3A). Expression continued to increase at 48 hours (~4 fold) in TBT and ROSI groups while expression in the LG group plateaued compared to controls (Figure 3A). Treatment with a selective PPARγ antagonist (T0070907) revealed that TBT-induced expression of miR-223 was strongly attenuated by antagonizing PPARγ (Figure 3B). In contrast, treatment with the RXR antagonist HX531 decreased miR-223 expression (Figure 3C) only marginally while treatment with a different RXR antagonist, UVI3003 (Figure 3D), had no effect (Figure 3D). Treatment with the RXR activator LG100268 induced expression of miR-223 modestly. Since PPARγ antagonists blunted miR-223 induction by ROSI and TBT, but RXR antagonists had no significant effect, we infer that TBT induced miR-223 expression by activating PPARγ rather than RXR.

**Figure 3.**
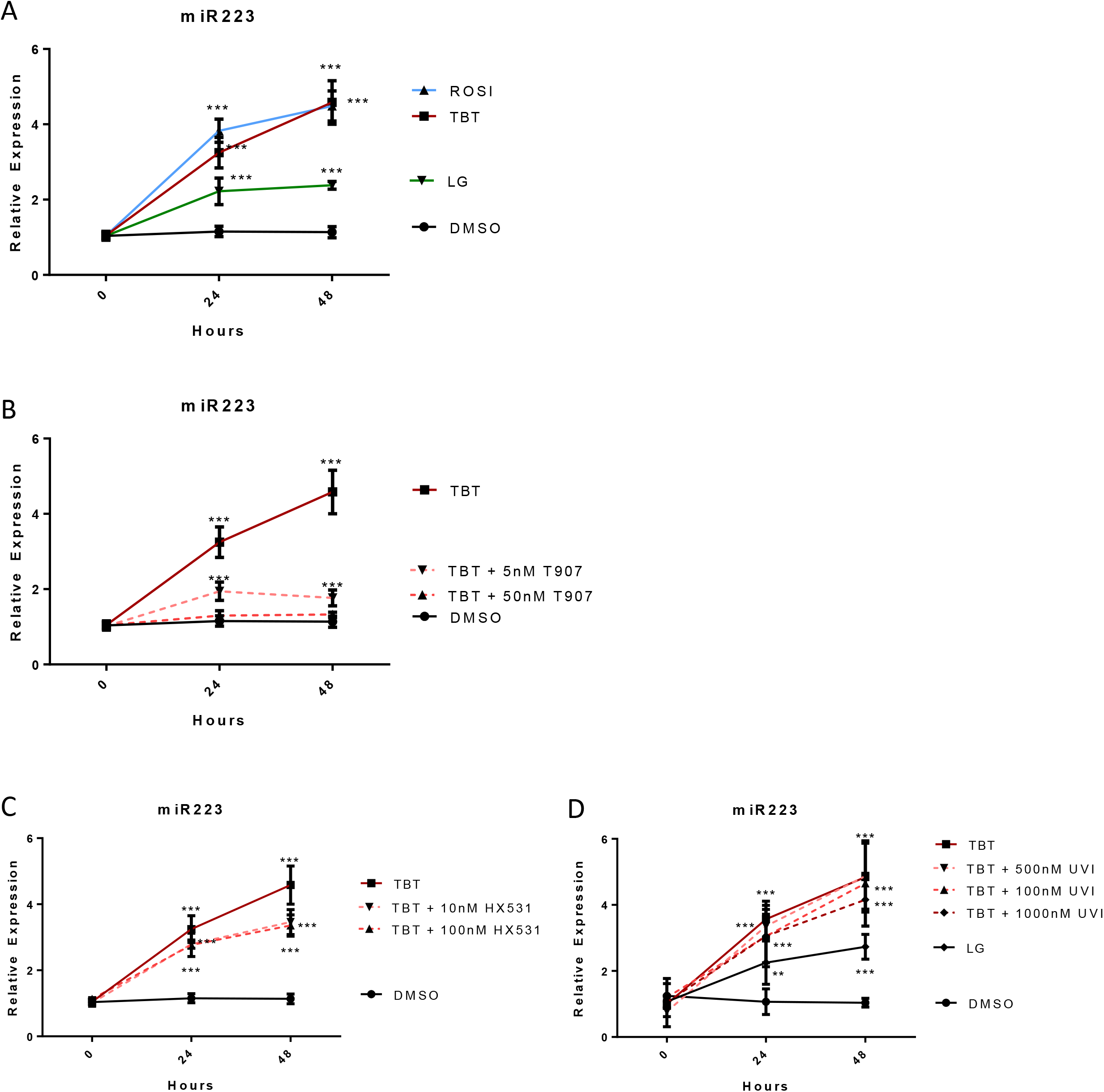
Effects of chemical inhibitors of nuclear receptors on TBT-induced miR223 expression in MSCs: qRT-PCR determination. (A) miR-223 expression under 0-48 hour exposure to 100 nM ROSI, 50 nM TBT, or 100nM LG268. (B) Expression under 0-48 hour expospure to 50 nM TBT in the presence of 5 or 50 nM T0070907. (C) Expression under 0-48 hour exposure to 50 nM TBT in the presence of 10 or 100 nM HX531. (D) Expression under 0-48-hour exposure to 50 nM TBT in presence of 100-1000 nM UVI3003. Expression was normalized to sno202 and presented as relative expression to the day 0 controls (mean ± SEM, **P < 0.01 and ***P < 0.001).

### TBT induces the binding of PPARγ to regulatory elements upstream of the pre-miR-223 coding region

PPARγ regulatory elements (PPREs) locating upstream of the pre-miR −223 coding region were previously described [28, 39]. We used chromatin immunoprecipitation (ChIP) with antibodies against PPARγ to identify which, if any, of these PPREs interacted with PPARγ upon exposure to TBT. Cells were treated for 48-hours with 50 nM TBT, 100 nM ROSI, or 100 nM LG268 in the presence or absence of T0070907 or HX531 in the TBT groups. Enrichment of PPREs was measured by qPCR with primer pairs flanking the PPREs. Compared with the negative control of IgG pull-down groups (Figure S4), our results showed no PPARγ binding enrichment in the PPRE −270 to −275 closest to the transcription start site of miR223 pre-RNA (Figure 4A). We found mild, but significant, enrichments in two PPREs, −1125 to −1130 and −1305 to −1310 (Figure 4C and 4D), and strong enrichment in two PPREs, −1011 to −1016 and −3919 to −3924 (Figure 4B and 4E). T0070907 treatment strongly decreased PPARγ enrichments of the four positive PPREs (Figure 4B-D) whereas HX531 weakly suppressed TBT induced PPARγ enrichments of three PPREs, −1011 to −1016, −1125 to −1130, and −1305 to −1310 (Figure 4B-D). Taken together, these ChIP assays demonstrated that TBT induced the binding of PPARγ to four out of five upstream PPREs of pre-miR-223, suggesting direct transcriptional activation of miR-223 expression by TBT-activated PPARγ.

**Figure 4.**
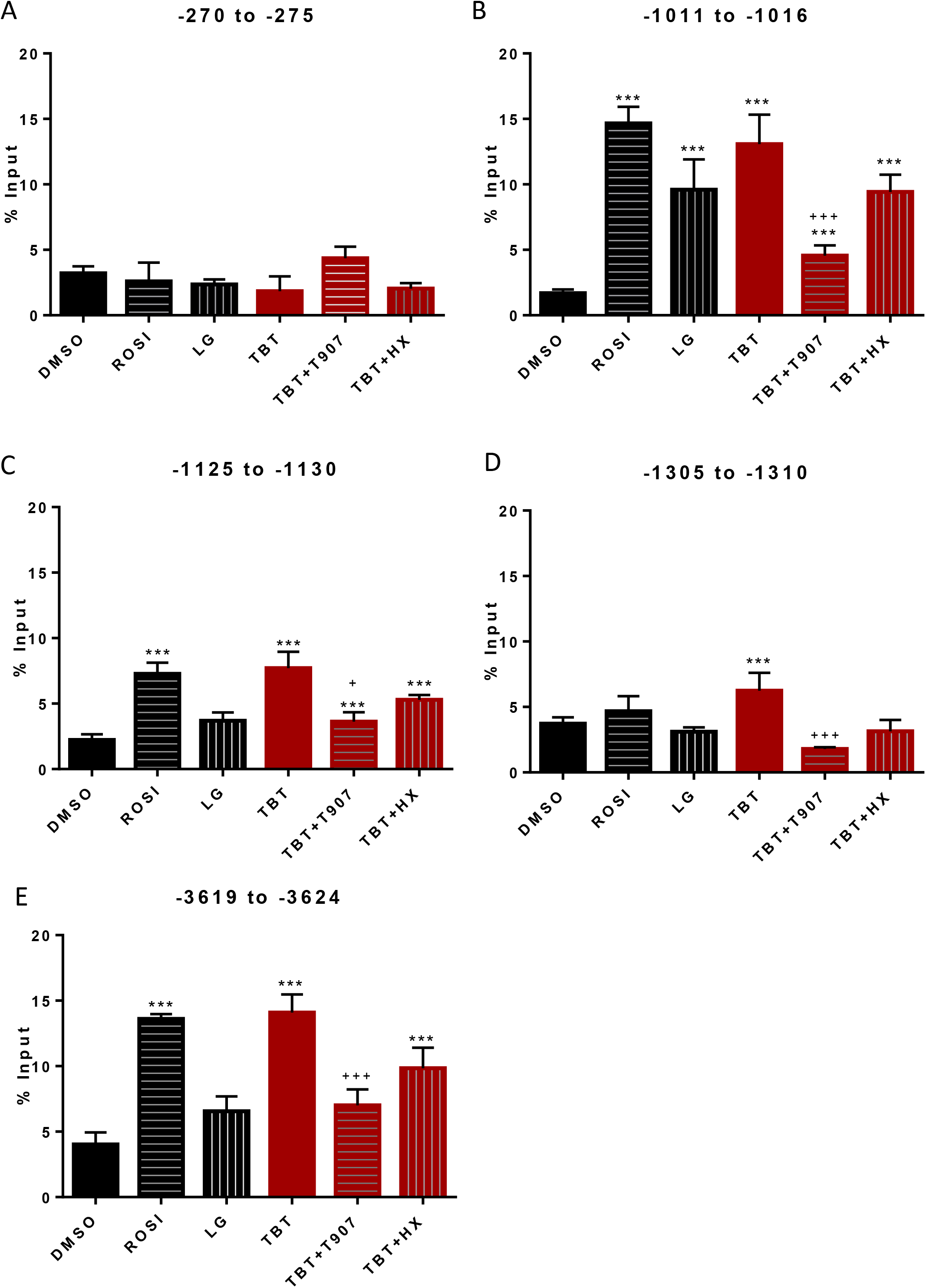
TBT-dependent binding of PPARγ to PPREs upstream of the pre-miR-223 transcription initiation site. MSCs were exposed to 100 nM ROSI, 50 nM TBT, 100nM LG268, or 50 nM TBT for 48 hours in presence of 50 nM T0070907 or 100 nM HX531. PCR-based ChIP assays were performed using an anti-PPARγ antibody for PPREs at the regions indicated in each panel. Each column represents mean ± SEM of 4 replicated assays (**P < 0.01 and ***P < 0.001 compared to control; + P < 0.05, ++ P < 0.01 and +++ P < 0.001 compared to TBT).

### miR223 expression in gonadal white adipose tissue of F2 and F3 male mice

The effects of prenatal TBT treatment on obesity have been reported as transgenerational and detectable in the F1, F2, F3, and F4 descendants of F0 mice exposed during pregnancy [9] or during pregnancy and lactation [17], whereas ROSI was unable to elicit transgenerational obesity [9]. The mechanism through which TBT acts to promote these transgenerational effects remains unclear. We next sought to examine the miR-223 levels in the adipose tissues of F2 and F3 animals ancestrally exposed to TBT following the procedure we previously published [17]. MicroRNA qPCR revealed a statistically significant increase in miR-223 expression in F2 and F3 adipose tissue of male mice exposed to regular chow diet (Figure 5A). Male offspring in the TBT group that were exposed to a diet with elevated fat levels (21.6% vs. 13.2% in chow) resulted in diet-induced obesity; these TBT group males showed strongly elevated miR-223 expression in both F2 (3.1 fold) and F3 (2.2 fold) generations compared with the vehicle group (Figure 5B). miR-223 levels were slightly increased in females from the TBT group vs controls in both chow diet and higher fat diet females, but this increase did not reach statistical significance (Figure 5C-D). These data demonstrate that the transgenerational obesity caused by ancestral exposure to TBT is associated with increased expression of miR-223 in white adipose tissue.

**Figure 5.**
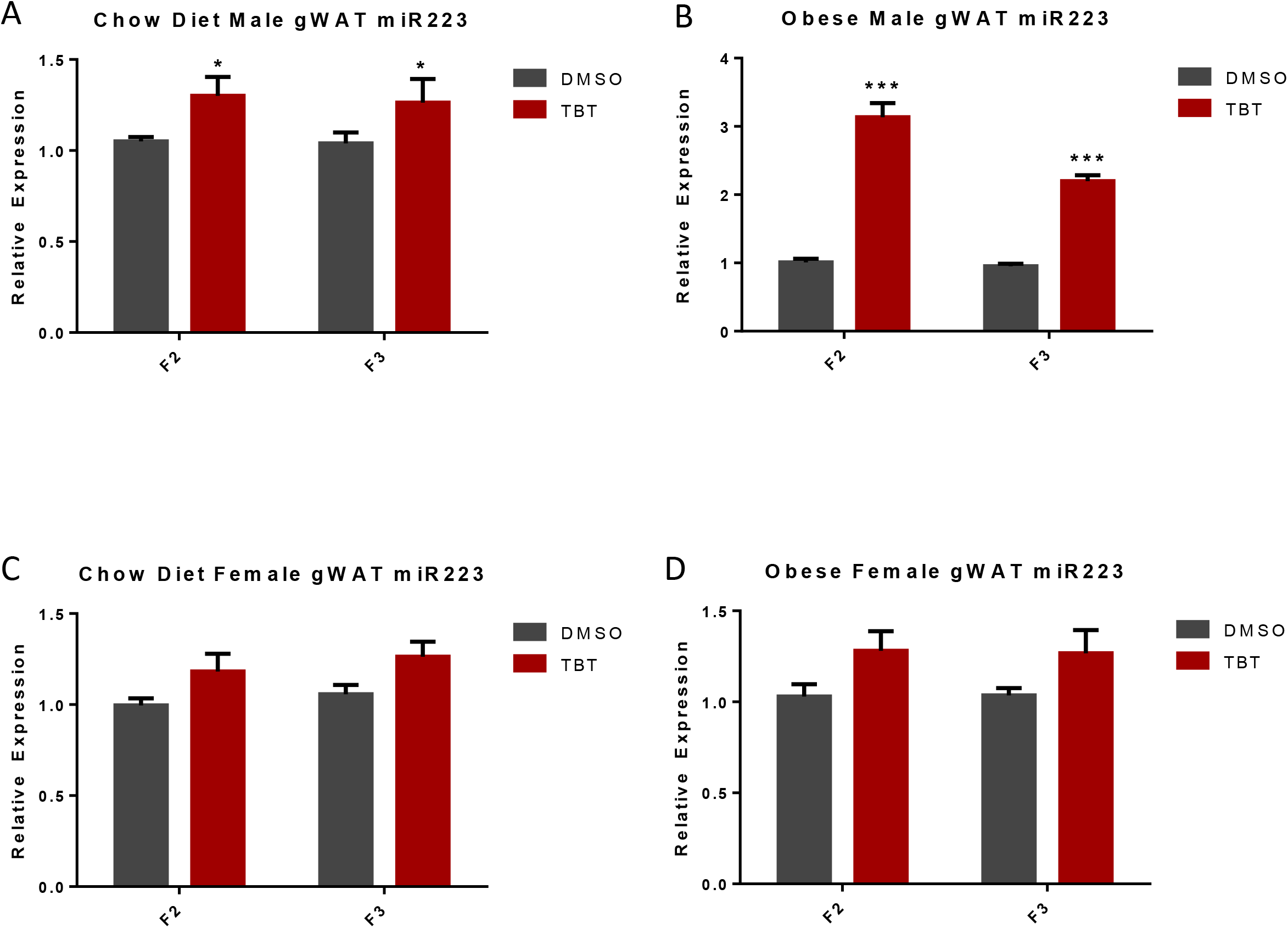
miR223 expression in gonadal white adipose tissue (gWAT) of F2 and F3 mice. Amounts of miR-223 mRNA in gWAT of males (A, B) and females (C, D) were determined using qRT-PCR. gWATs were collected before (A, C) or after (B, D) the high-fat diet challenge. Expression was normalized to sno202 and presented as relative expression to DMSO controls (mean ± SEM, n = 11; *P < 0.05, **P < 0.01 and ***P < 0.001).

## Discussion

We previously examined effects of TBT on adipocytic differentiation of mouse MSCs and analyzed the transcriptomal profiles during the course of their in vitro adipogenesis [13]. Here, we utilized these transcriptomal data to identify microRNAs that could be involved in adipogenesis in MSCs. We revealed roles of microRNAs in TBT-induced adipogenesis, particularly miR-223. Inhibition of PPARγ action by specific antagonists or microRNA suppression assays using miR-neutralizing antisense RNAs identified a critical role for miR-223 in adipogenesis. Our finding that PPARγ binds to PPREs in the promoter of the gene encoding pre-miR-223 in an agonist-dependent manner supports a direct role of PPARγ in regulating miR-223 expression.

To date, knowledge about miR-223 in obesity has been limited to its role in adipose-resident macrophages [27] as a regulator of myeloid lineage development [23, 40, 41]. Here, we identified an axis involving TBT, PPARγ, and miR-223 that enhances differentiation of white adipocytes from MSCs. Overexpression of synthetic miR-223 mimic or lentivirally-expressed miR-223 was reported to potentiate adipocyte lipid accumulation by enhancing the expression of PPARγ, C/EBPα, adipsin, and aP2 in both ST2 and C3H10T_1/2_ cell models [29]. Conversely, knockdown of the TBT-induced miR-223 using a specific antisense RNA in our study blunted lipid accumulation in adipocytes. In addition, the three miR-223 target genes we measured were reported to play important roles in adipocytes. FOXP1 overexpression impaired adaptive thermogenesis in adipocytes and led to diet-induced obesity [42]. ZEB1 was shown to be a critical transcription factor regulating adipose tissue accumulation in C57BL/6 mice [43], A point mutation in ZEB1 also increased body weight in Twirler mice model [44]. NFIB is a sequence-specific DNA binding protein that interacts with KDM4D to promote the expression of adipogenic genes [45]. These findings support a model in which TBT-activates PPARγ, inducing miR-223 expression and adipocyte differentiation from multipotent MSCs.

Regulation of miR-223 expression has been studied by analysis of cis-acting elements within the proximal promoter. Several potential binding sites for c/EBPs were identified in the miR-223 promoter. In human myeloid precursors, expression of miR-223 was reported to be upregulated by c/EBPs [46], contributing to granulocyte differentiation [47]. Studies in human endothelial cells [48] and murine macrophages [49] also suggested that c/EBPs could induce miR-223 expression. Taken together, these studies identified binding of c/EBPs to the proximal promoter of miR-223 as an important transcriptional regulator of miR-223 expression.

Based on our previous findings that TBT bound to both members of the PPARγ-RXR heterodimer to modulate adipogenesis [12, 13], we sought to elucidate molecular mechanisms through which TBT promoted adipogenesis via regulation of miR-223 in MSCs. We examined PPREs known to be important for the expression of pre-miR-223 [28] and identified four PPREs to which TBT-activated PPARγ bound in an agonist-dependent manner. These data show that TBT-activated PPARγ up-regulated expression of pre-miR-223 via binding to the PPREs in its promoter.

We previously observed that prenatal exposure of pregnant F0 mouse dams to TBT led to heritable effects including increased WAT depot sizes, reprogramming of MSC fate toward the adipose lineage, and increased hepatic lipid storage in F1, F2 and F3 generations without further exposure [9]. The observed transgenerational predisposition to obesity was strongly male-specific in all three generations [9, 16, 17]. Here we identified a mechanism through which TBT activated PPARγ, upregulating expression of mir-223 leading to post-transcriptional regulation of important target genes in adipogenesis. miR-223 levels remained higher in F2 and F3 male descendants of F0 dams exposed to TBT throughout pregnancy and lactation [17]. Elevated miR-223 levels preceded the development of diet-induced obesity in the F2 and F3 males; miR-223 levels were further increased by HFD exposure. Future studies should address the molecular mechanisms underlying the transgenerationally affected expression of miR-223. One possibility suggested by our previous studies is altered higher-order chromatin structure [17, 50]. Whether the transgenerationally altered miR-223 expression is part of the vehicle of non-genetic inheritance or a consequence of it, our study demonstrates convincingly that obesogen-induced miR-223 facilitates the increased differentiation of MSCs into white adipocytes. While it has been proposed that obesogen-induced changes in expression of non-coding RNAs can play a role in obesity [51], our current study provides the first evidence that this is indeed the case.

## Acknowledgements

Funded by grants from the NIH (ES023316, ES031139) to B.B. and T.S. and the Environmental Protection Agency (STAR FP917800 to B.M.S.).

---

**S1.**
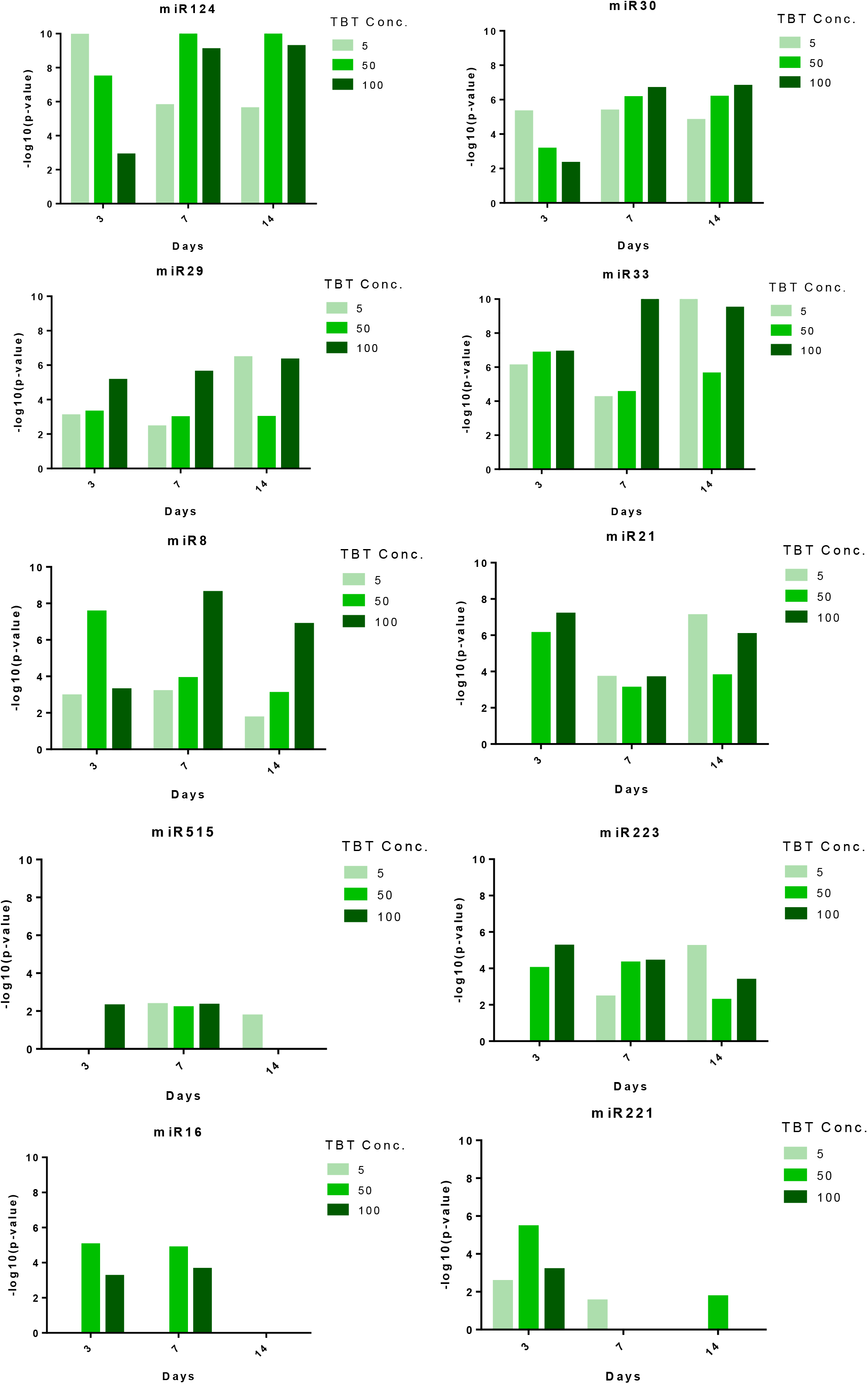

**S2.**
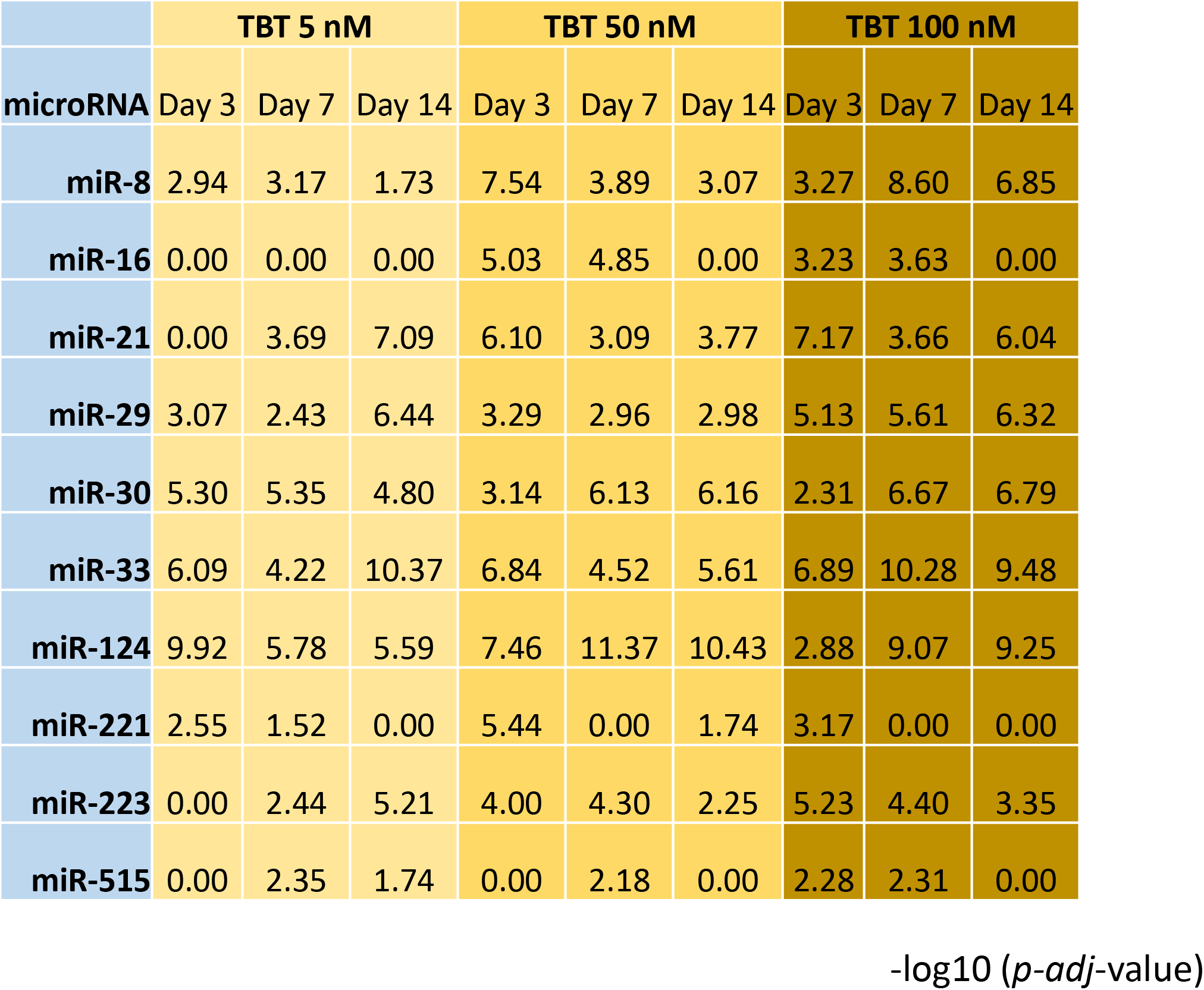

**S3.**
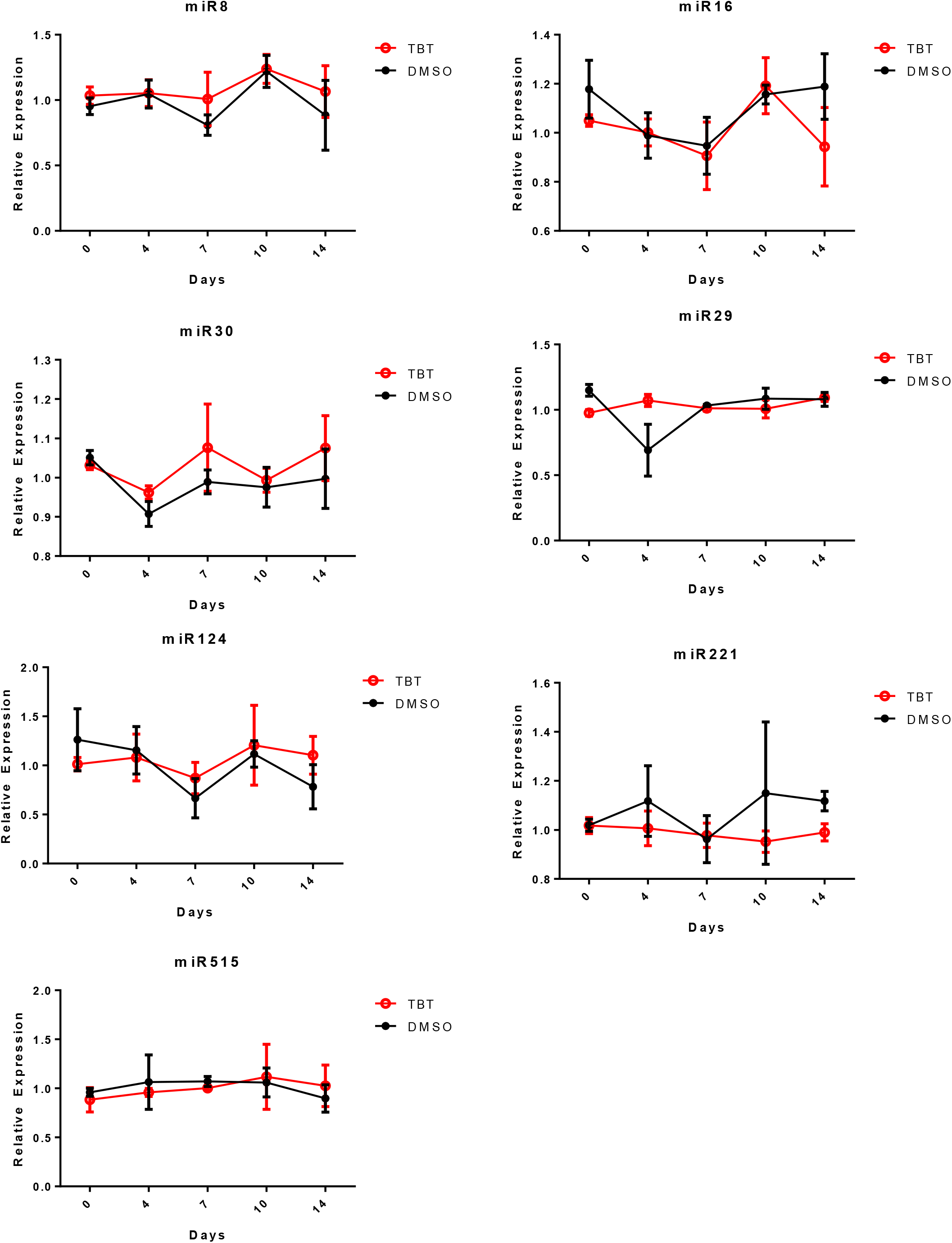

**S4.**
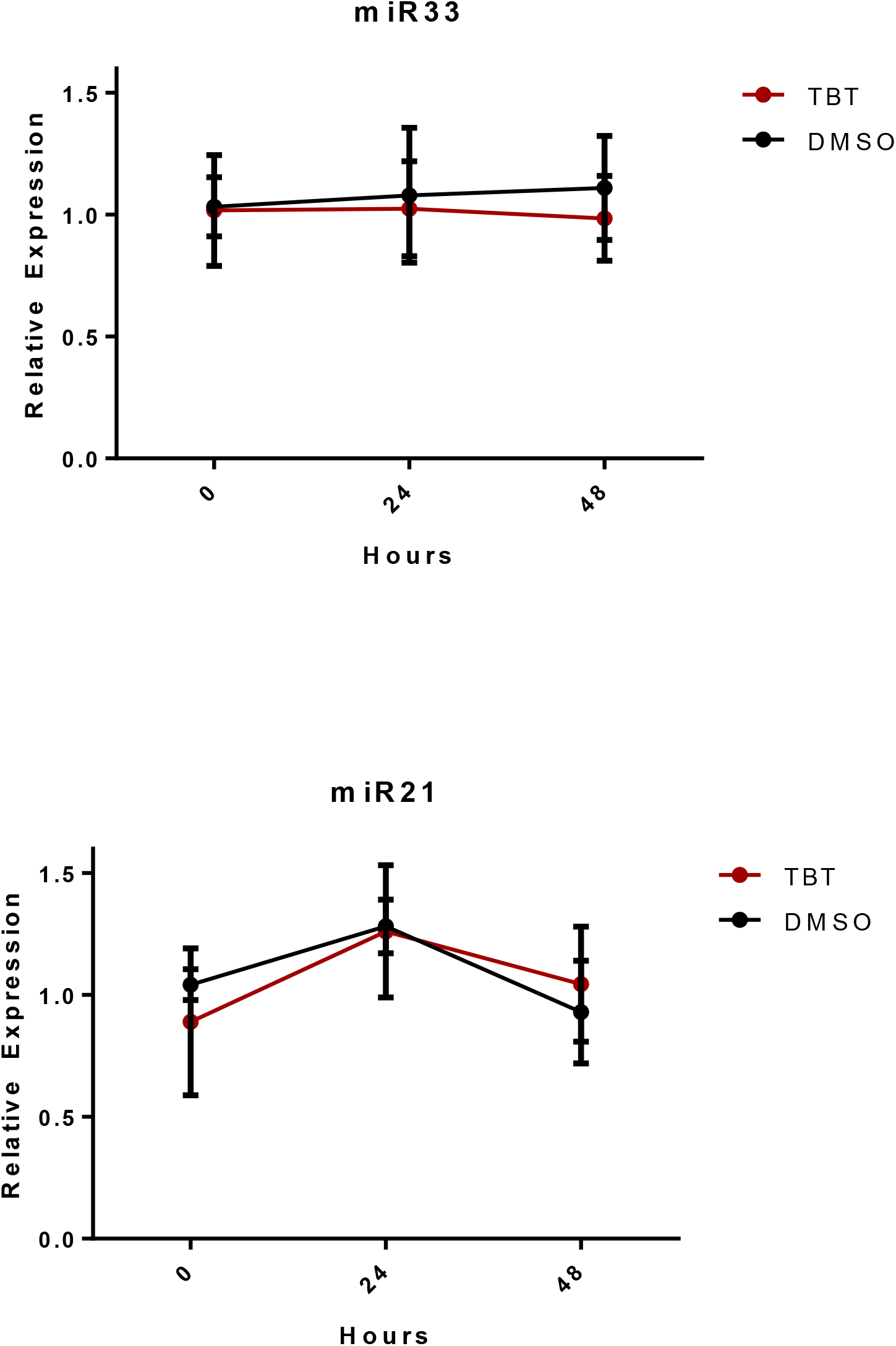

**S5.**
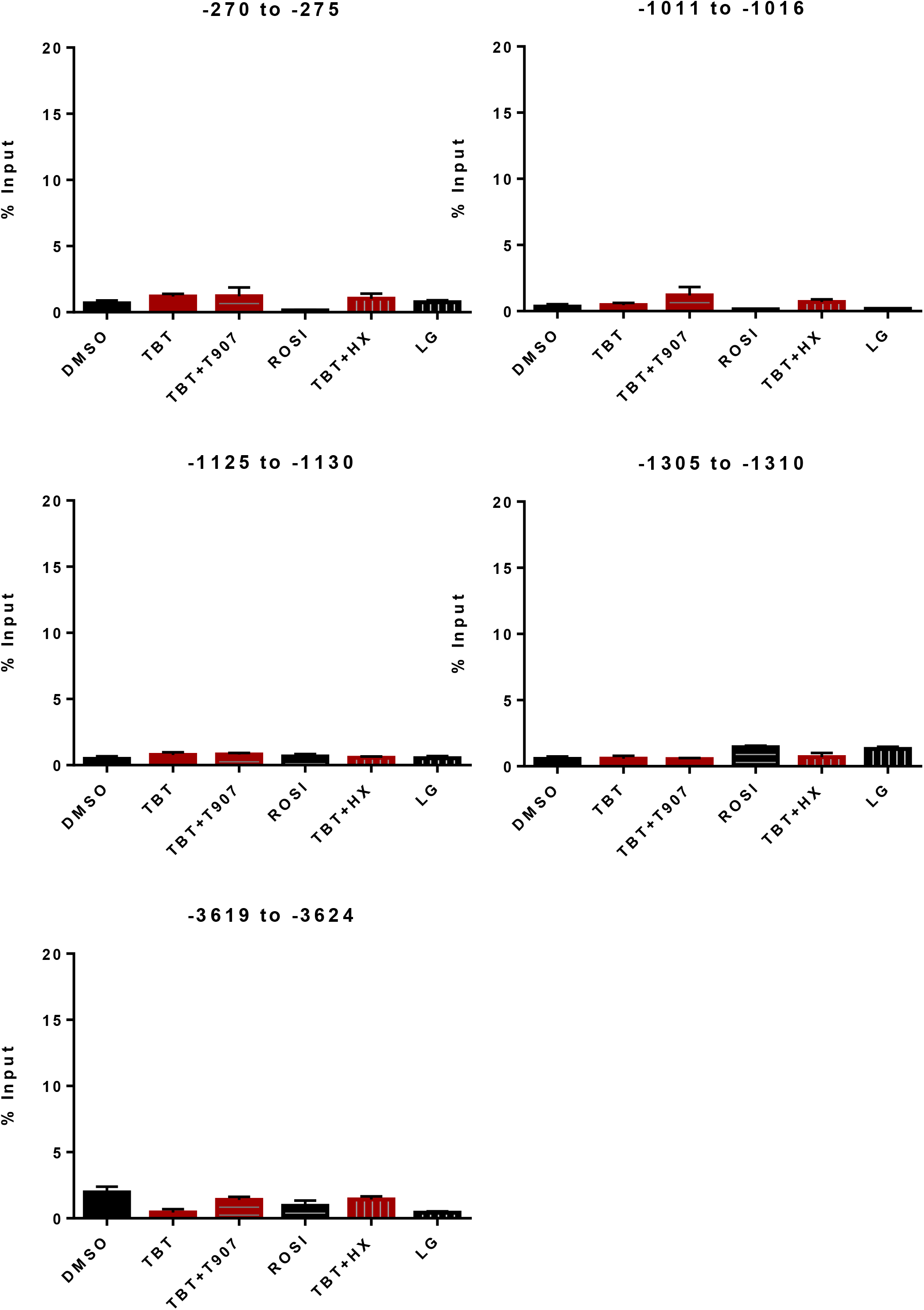

